# Zinc-dependent Nucleosome Reorganization by PARP2

**DOI:** 10.1101/2023.10.17.562808

**Authors:** Natalya Maluchenko, Alexandra Saulina, Olga Geraskina, Elena Kotova, Anna Korovina, Alexey Feofanov, Vasily Studitsky

## Abstract

Poly(ADP-ribose)polymerase 2 (PARP2) is a nuclear protein that acts as a DNA damage sensor; it recruits the repair enzymes to a DNA damage site and facilitates formation of the repair complex. Using single particle Förster resonance energy transfer microscopy and electrophoretic mobility shift assay (EMSA) we demonstrated that PARP2 forms complexes with a nucleosome containing different number of PARP2 molecules without altering conformation of nucleosomal DNA both in the presence and in the absence of Mg^2+^or Ca^2+^ions. In contrast, Zn^2+^ ions directly interact with PARP2 inducing a local alteration of the secondary structure of the protein and PARP2-mediated, reversible structural reorganization of nucleosomal DNA. AutoPARylation activity of PARP2 is enhanced by Mg^2+^ ions and modulated by Zn^2+^ ions: suppressed or enhanced depending on the occupancy of two functionally different Zn^2+^ binding sites. The data suggest that Zn^2+^/PARP2-induced nucleosome reorganization and transient changes in the concentration of the cations could modulate PARP2 activity and the DNA damage response.

**Significance Statement:** PARP2 recognizes and binds DNA damage sites, recruits the repair enzymes to these sites and facilitates formation of the repair complex. Zn^2+^-induced structural reorganization of nucleosomal DNA in the complex with PARP2, which is reported in the paper, could modulate the DNA damage response. The obtained data indicate the existence of specific binding sites of Mg^2+^ and Zn^2+^ ions in and/or near the catalytic domain of PARP2, which modulate strongly, differently and ion-specifically PARylation activity of PARP2, which is important for maintaining genome stability, adaptation of cells to stress, regulation of gene expression and antioxidant defense.

## Introduction

PARP2 is a nuclear enzyme and a member of the poly(ADP-ribose)polymerase (PARP) family of proteins. PARP2 has a multi-domain structure, which includes N-terminal region (NTR), tryptophan-glycine–arginine-rich (WGR), α-helical and catalytic domains (1). PARP2 is involved in various differentiation processes including spermatogenesis (2), adipogenesis (3), thymocyte survival (4) and endometrial receptivity (5). It plays a special role in maintaining genome stability, acting as a damage sensor that recruits DNA repair enzymes including base excision repair (BER) factors, X-ray repair cross complementing 1 (XRCC1), DNA (apurinic/apyrimidinic site) endonuclease 1 (APE1), DNA polymerase β (Polβ) and DNA ligase (6). PARP2 catalyzes the polyADP-ribosylation (PARylation) of proteins including PARP2 itself and histones (7) and provides the synthesis of 15–25% of the poly(ADP-ribose)polymer (PAR) in cells (8). PARP2 triggers reactions that promote cell adaptation to stress (9). Under genotoxic stress, PARP2 binds to RNA in the nucleoli, thus participating in a complex network targeting RNA preservation during the cellular response to DNA damage (10). PARP2 is able to partially compensate for the DNA repair functions of PARP1 after knockout of PARP1, while a double knockout of PARP1 and PARP2 is lethal (11). PARP2 can also regulate PARP1-catalyzed PAR synthesis (6, 12)

The molecular mechanisms of PARP2 action in chromatin are realized *via* its binding to DNA. The substrates recognized by PARP2 include DNA containing single and double stranded breaks, flaps and overhangs, hairpins, gaps of various lengths, sticky ends as well as branched DNA with junctions of 4 or 3 strands (6, 12, 13). PARP2 was proposed to act at later stages of DNA repair as compared to PARP1 (1, 6, 12, 14).

Recent studies have shown that PARP2 uses the WGR domain, in addition to NTR for DNA recognition (1, 15-17). WGR plays a key role in PARP2 binding to DNA and subsequent DNA-dependent activation of PARP2 catalytic activity (17, 18). WGR binding to DNA destabilizes the α-helical regulatory domain and allows the catalytic domain to bind the substrate (NAD+), carry out PARylation and, finally, recruit DNA-repair proteins (16). Since genomic DNA exists in a cell in the form of chromatin, nucleosomes are inevitably involved in the interaction with PARP2. Recent studies demonstrate that the affinity of PARP2 for nucleosomal DNA is higher than for naked DNA (13), and WGR domain is responsible for the binding to nucleosomal DNA (19, 20).

Here we report that, in spite of the apparent absence of the specialized Zn^2+^-binding domains (e.g. zinc fingers), interaction of PARP2 with nucleosomes is modulated by Zn^2+^ ions. Furthermore, PARP2 binding to a nucleosome in the presence of Zn^2+^ ions is accompanied by changes in the structure of nucleosomal DNA. This is not observed in the presence of Ca^2+^ or Mg^2+^ ions. PARylation activity of PARP2 is shown to be considerably enhanced by Mg^2+^ ions. Zn^2+^ ions suppress or enhance it depending on the occupancy of two functionally different Zn^2+^ binding sites. Zn^2+^ ions bind to PARP2 and induce subtle changes in the secondary structure of the enzyme. Localization of the Zn^2+^ binding site as well as Zn^2+^-dependent features of PARP2 functioning are discussed.

## Materials and Methods

### Reagents

The following reagents were used: EDTA (Amresco, USA), ZnCl_2_ (Fluka, Switzerland), CaCl_2_, MgCl_2_, NAD+, sodium dodecyl sulfate (SDS), dithiothreitol (DTT), polyvinylidene fluoride membrane (Sigma Aldrich, USA), mouse monoclonal antibodies against PAR (clone 10H, Tulip BioLabs, USA), anti-mouse antibodies conjugated with horseradish peroxidase (Bio-Rad, Hercules, CA, USA)

### Proteins, DNA templates and nucleosomes

Recombinant mouse full-length PARP2 was obtained in the baculovirus expression system using Sf9 insect cells as described earlier (21) and was kindly provided by Prof. O. Lavrik.

Fluorescently-labelled 187-bp DNA templates were obtained by polymerase chain reaction (PCR) using the plasmid containing nucleosome-positioning sequence s603-42A and fluorescently labeled oligonucleotides as primers. The following oligonucleotides (Lumiprobe, Russia) were used:

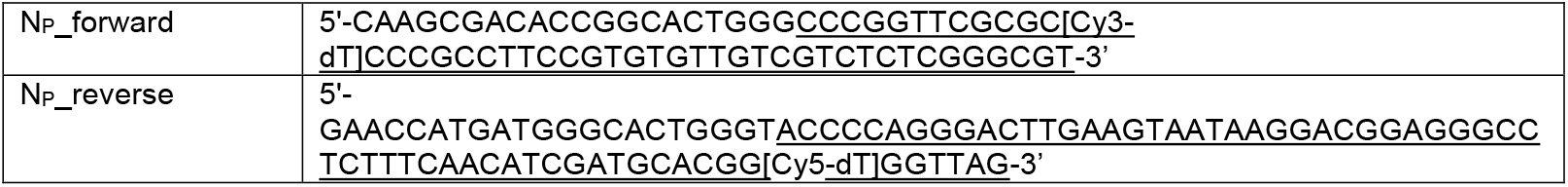
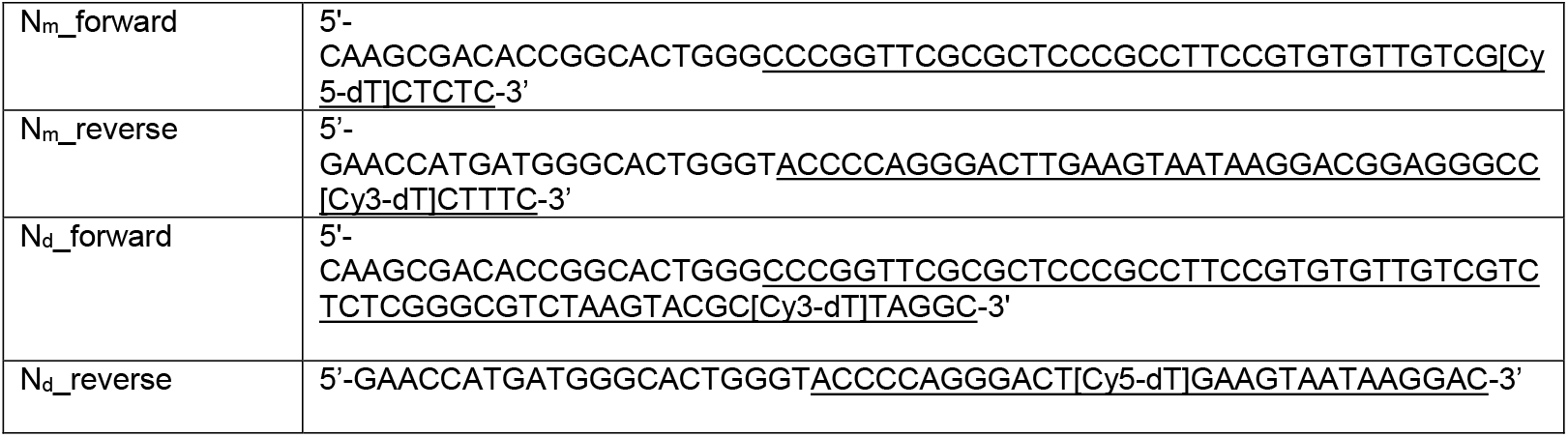

PCR products were purified from 2% agarose gel and extracted by QIAquick Gel Extraction Kit (Qiagen) following the manufacturer’s protocol.

Nucleosomes were assembled using fluorescently-labelled DNA templates and chicken donor chromatin without linker histone H1 as described earlier (22). Assembled nucleosomes were purified by the electrophoresis in the 4.5% polyacrylamide gel (acrylamide:bisacrylamide 39:1; 0.5× TBE buffer, pH 8.0), isolated from the gel by extraction in a buffer containing 10 mM HEPES-NaOH (pH 8.0), 0.2 mM EDTA, 0.2 mg/ml bovine serum albumin and stored at 4 °C. The purified nucleosomes contained less than 3% of histone-free DNA.

### spFRET experiments in solution

In the study of nucleosome-PARP2 interactions, nucleosomes (1-2 nM) were incubated with PARP2 (12.5 - 200 nM) in buffer A (50 mM Tris-HCl pH 7.5; 40 mM NaCl; 1 mM β-mercaptoethanol (β-ME); 0.1% NP40) at 25 °C for 30 min. When indicated, buffer A was supplemented with 0.2 - 3 mM ZnCl_2_ or 5 mM CaCl_2_ or MgCl_2_. To study the stability of Zn^2+^-PARP2-nucleosome complexes, EDTA (10 mM) was added to PARP2-nucleosome complexes preformed for 30 min in buffer A containing 200 nM PARP2 and 0.3 mM Zn^2+^ ions and incubated for another 15 min after adding EDTA.

In the studies of PARylation, NAD+ (0.5-10 μM) was mixed with nucleosomes (1-2 nM) and PARP2 (100 nM) in buffer A and incubated for 45 min at 25° C.

spFRET measurements from freely diffusing nucleosomes in solution were performed as described previously (23). Each measured single nucleosome was characterized by FRET between Cy3 and Cy5 labels calculated as a proximity ratio (E_PR_):

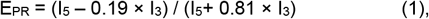

Where I_3_ and I_5_ are fluorescence intensities of Cy3 and Cy5, respectively, and coefficients 0.19 and 0.81 provide correction for the spectral cross-talk between Cy3 and Cy5 detection channels. E_PR_ is a FRET efficiency not corrected for quantum yields of labels and an instrumentation factor. Relative frequency distributions of nucleosomes by E_PR_ values (2000-5000 nucleosomes per experiment; 3 independent experiments) were plotted and further analyzed as a superposition of several normal (Gaussian) distributions for free nucleosomes or nucleosome-PARP2 complexes.

### EMSA (Electrophoretic Mobility Shift Assay) experiments

Sample preparation for EMSA was the same as for spFRET microscopy, except for slightly higher concentration of nucleosomes (2-3 nM). The PARP2-nucleosome complexes were subjected to electrophoresis in 5% polyacrylamide gel (acrylamide:bisacrylamide 59:1; 0.2×TBE buffer). Electrophoresis was performed under native conditions at 140 V, + 4°C for ∽1.5 h. The gels were scanned using Amersham Typhoon RGB imager (Cytiva, Sweden). Fluorescence was excited in the gel at the 532 nm wavelength and recorded in the 570–610 nm (Cy3 signal) and 650–700 nm (FRET signal of Cy5) spectral regions. The resulting images were encoded by green (Cy3 signal) and red (FRET signal of Cy5) colors and merged.

### Single particle fluorescence intensity analysis of nucleosomes in the gel

For the analysis of stoichiometry of the PARP2-nucleosome complexes, gels obtained in EMSA experiments were placed between object and cover glasses and subjected to single particle fluorescence intensity analysis as described previously (24). Measurements were performed using the 633 nm excitation wavelength and the 650– 800 nm detection range, thus exciting and detecting fluorescence of Cy5 only. This approach made it possible to estimate the number of nucleosomes per single particle (PARP2-nucleosome complex) for each gel lane and make conclusion about the stoichiometry PARP2-nucleosome complexes separated in gel.

### Western Blots (WB)

Nucleosomes (3 nM) were preincubated with PARP2 (100 nM) in the buffer 50 mM Tris-HCl, (pH 7.5), 40 mM NaCl, 0.1% NP40 (supplemented with 5 mM MgCl_2_ or 0.3 mM ZnCl_2_ when indicated) for 30 min followed by 5 min incubation with 10 mM EDTA (when indicated) and 30 min incubation with NAD+ (0, 1, 2.5 or 5 µM) in the same buffer. Probes were subjected to electrophoresis in 4-12% bis-Tris gradient gel in the NuPAGE™ MES SDS Running Buffer (50 mM MES, 50 mM Tris-HCl, 0.1% SDS, 1 mM EDTA, pH 7.3) at 130 V. Protein transfer on polyvinylidene fluoride membrane was performed in the transfer buffer (25 mM Bicine, 25 mM Bis-Tris (free base), 1 mM EDTA pH 7.2) with 20% methanol at 4°C and 70 V for 1 h. The membrane was incubated for 60 min in the PBS-T solution (2.7 mM KCl, 8 mM Na_2_HPO_4_, 2 mM KH_2_PO_4_, 37 mM NaCl, 0.5% Tween 20) supplemented with the 5% skimmed milk (prepared from dry powder) and washed with the 1×PBS-T solution for 5 min. Then the membrane was incubated with mouse monoclonal antibodies against PAR for 60 min in PBS-T/5% milk followed by incubation with the secondary anti-mouse antibodies conjugated with horseradish peroxidase (Bio-Rad, Hercules, CA, USA) for 60 min in the PBS-T/5% milk solution. The washing procedure was carried out after each step of incubation. Immunodetection was performed using the Super Signal West Pico Chemiluminescent Substrate (Thermo Fisher Scientific, Waltham, MA, USA) for 3 min.

### Circular dichroism (CD) spectroscopy

The samples, which contained 0.5 g/L PARP2 in buffer (12.5 mM NaCl, 25 mM Tris-HCl pH 7.5, 0.5 mM DTT, 12.5% glycerol) with or without 0.3 mM ZnCl_2_, were prepared immediately after defrosting the concentrated solution of PARP2 and stored on ice for no longer than 30 min before the measurement.

CD spectra of PARP-2 were recorded in the 190-250 nm spectral range with 0.2 nm step using Jasco-810 spectrophotometer (Jasco, Japan) and the quartz SUPRASIL® cuvette of 0.1 mm thickness (Hellma, Germany). The blank subtraction was applied to all spectra, and the data of two independent experiments were averaged.

Secondary structure analysis was performed by CDPro package programs using the SP43 reference set. The content of canonical secondary structures of various types predicted by Continll, SELCON 3 and CDSSTR programs were averaged to improve the reliability of the analysis (25). Statistical significance of the secondary structure difference was determined using the Holm-Sidak method.

## Results

### Experimental approach

Interaction of nucleosomes with PARP2 was studied by combining the electrophoretic mobility shift assay (EMSA), fluorescent microscopy of single particles/complexes based on Förster resonance energy transfer (spFRET microscopy), Western blotting (WB) and circular dichroism (CD) spectroscopy.

To support spFRET measurements (and, in part, EMSA), the nucleosomes were labeled with a donor-acceptor pair of Cy3 and Cy5 fluorophores (Figure 1a) attached to the neighboring gyres of nucleosomal DNA at the following positions: 13 and 91 bp (N_p_ nucleosomes), 35 and 112 bp (N_m_ nucleosomes) or 57 and 135 bp (N_d_ nucleosomes) from the boundary of the 603 nucleosome positioning DNA sequence (23, 26). The labels placed at these positions allow efficient FRET and probing structural changes near and far from the boundary of the nucleosomal DNA (23, 26, 27). Single nucleosomes or their complexes with PARP2 freely diffusing in a solution were subjected to spFRET measurements and the data were presented as graphs describing the frequency distributions of nucleosomes by proximity ratio E_PR_ (Figure 1b, d). E_PR_, which was calculated for each measured nucleosome, is the analog of the FRET efficiency without correction for the instrumental factor and fluorescence quantum yields of the Cy3 and Cy5 labels.

**Figure 1.**
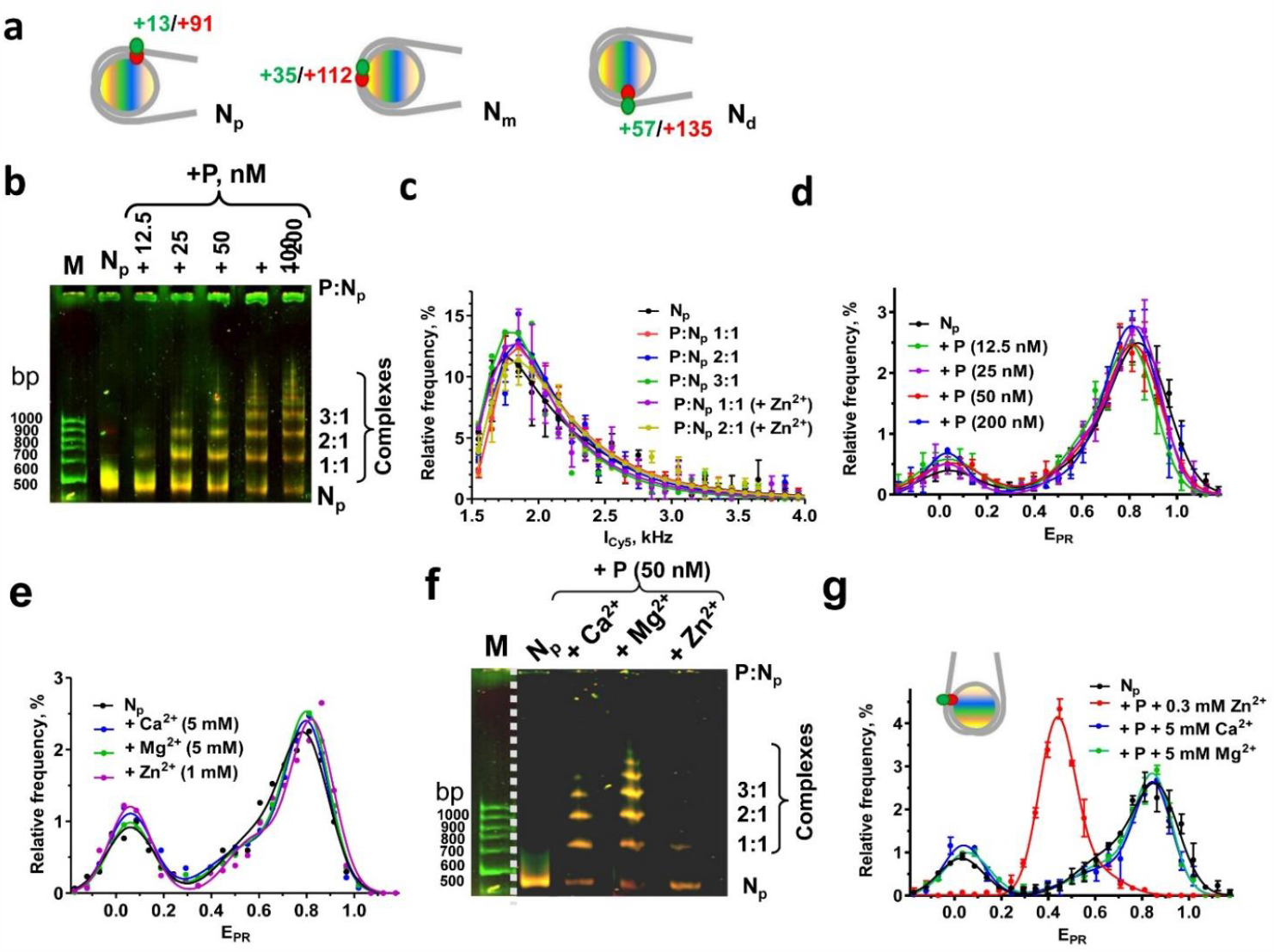
Multiple PARP2 molecules can bind to a nucleosome and induce Zn^2+^-dependent nucleosome reorganization. **a)** Positions of Cy3 (green) and Cy5 (red) labels in the nucleosomes. Numbers indicate distances (bp) from the boundary of nucleosomal DNA to the labeled nucleotides. **b)** Analysis of N_P_ nucleosomes and their complexes with PARP2 (P, 12.5-200 nM) formed in buffer A by non-denaturing PAGE. P: N_p_ is the proposed stoichiometry of the PARP2 complexes with nucleosomes observed in the gel. M – DNA markers. **c)** Frequency distributions (E^PR^ profiles) of N_P_ nucleosomes and their complexes with PARP2 measured by single particle fluorescence (spFRET) microscopy within different bands of the gels shown in **Figure 1b, f**. **d)** E_PR_ profiles of N_P_ nucleosomes measured with spFRET microscopy in buffer A in the absence and presence of PARP2 (P). Concentration of PARP2 varied from 12.5 to 200 nM; **e)** E_PR_ profiles of N_P_ nucleosomes in buffer A in the absence of divalent cations (N_p_) or in the presence of Zn^2+^ (1 mM), Ca^2+^ (5 mM) or Mg^2+^ (5 mM) ions. **f)** Analysis of N_P_ nucleosomes and their complexes with PARP2 (P, 50 nM) formed in buffer A in the absence of divalent cations (N_p_) or in the presence of Zn^2+^ (0.3 mM), Ca^2+^ (5 mM) or Mg^2+^ (5 mM) ions by non-denaturing PAGE. P: N_P_ is a proposed stoichiometry of the PARP2 complexes with nucleosomes observed in the gel. M – DNA markers. **g)** E_PR_ profiles of N_P_ nucleosomes in buffer A(N) and complexes of N_P_ with PARP2 (P, 50 nM) in buffer A supplemented with Zn^2+^ (0.3 mM), Ca^2+^ (5 mM) or Mg^2+^ (5 mM). **(c-e**,**g)** Statistics: mean ± SEM; 3 independent experiments; ∼3000 particles per experiment.

EMSA was used to analyze formation of complexes between nucleosomes and PARP2 under non-denaturing (native) conditions. Preservation of the nucleosome structure in the complexes was monitored using the FRET-in-gel analysis (28). Differences in PARylation activity of PARP2 in the complex with nucleosomes at different conditions were compared using WB. Structural changes in PARP2 were analyzed with CD spectroscopy.

### Complexes of nucleosomes with PARP2: EMSA and spFRET microscopy

According to EMSA, PARP2 forms complexes with nucleosomes in buffer A (50 mM Tris-HCl pH 7.5; 40 mM NaCl; 1 mM β-mercaptoethanol (β-ME); 0.1% NP40) at the concentration of the enzyme of ∼12.5 nM and higher (Figure 1b). A 50% decrease in the number of free nucleosomes is observed at 40±10 nM PARP2 according to the densitometry analysis of the gel image. An increase in PARP2 concentration is accompanied by formation of multiple complexes having different electrophoretic mobilities in the gel (Figure 1b). The color of the bands in gel can vary from green (no FRET) to yellow (high FRET) and finally to red (100% FRET) depending on the structure of the nucleosomes. The bands in the gel corresponding to free nucleosomes and different PARP2-nucleosome complexes have similar yellow color, suggesting that different complexes have similar structures of nucleosomal DNA (Figure 1b). Since formation of multiple complexes can indicate different stoichiometry of interacting components, we have analyzed fluorescence intensity of single complexes within particular bands in gel to evaluate whether the number of nucleosomes in the complexes with PARP2 varies. Fluorescence of Cy5 label was selectively excited and measured with single particle fluorescence microscopy. Comparison of frequency distributions within different gel bands corresponding to different complexes by fluorescence intensity, with the frequency distribution of intact nucleosomes in gel did not reveal significant differences (Figure 1c). The data suggest that different complexes observed in the gel contain one nucleosome per complex, and the different mobilities are explained by the binding of different number of PARP2 molecules to the nucleosome (Figure 1b).

Both the spFRET and EMSA data indicate absence of structural changes in nucleosomes upon formation of the complexes with PARP2 in buffer A: the E_PR_ profiles of nucleosomes are very similar in the absence and presence of PARP2 and reveal two subpopulations of nucleosomes in the solution (Figure 1d). Major subpopulation is characterized by higher E_PR_ values (distribution maximum at 0.8) and corresponds to nucleosomes, where neighboring gyres of nucleosomal are close to each other at the labeled sites. Minor subpopulation is characterized by much lower E_PR_ values (distribution maximum at 0.03) and corresponds to histone-free DNA and/or nucleosomes, in which the distance between DNA gyres increased considerably, for example, due to so-called nucleosome breathing (29). SpFRET analysis shows absence of structural changes in nucleosomes in the wide range of PARP2 concentrations (12.5-200 nM, Figure 1d), when various types of complexes of different stoichiometry are formed according to EMSA data (Figure 1b).

### Zn^2+^ selectively affects the structure of the nucleosome-PARP2 complexes

The nucleoplasm contains submillimolar concentrations of Ca2+, Mg2+ and Zn2+cations that could modulate the activity of nuclear proteins. Millimolar concentrations of Ca^2+^, Mg^2+^ or Zn^2+^ ions do not affect the nucleosome structure according to the spFRET data (Figure 1g). Furthermore, Ca^2+^ or Mg^2 +^ ions do not affect the structures of PARP2-nucleosome complexes revealed by EMSA and spFRET microscopy: 4-5 complexes with different stoichiometry are formed at the 50 nM concentration of PARP2 (Figure 1f); the structure of N^P^ nucleosomes is not changed in these complexes (Figure 1e, f). In contrast to Ca^2+^ and Mg^2+^ ions, the presence of Zn^2+^ ions induce alterations of PARP2-nucleosome complexes (Figure 1f, g). Analysis of EMSA and spFRET data shows that only two types of PARP2-nucleosome complexes are formed in the presence of Zn^2+^ ions (Figure 1f); both complexes contain one nucleosome (Figure 1c) and different numbers of PARP2 molecules per nucleosome (1:1 and 2:1); the structure of N_P_ nucleosomes is changed in these complexes (Figure 1f). In the E_PR_ profile, the changes in the nucleosome structure induced by combination of PARP2 and Zn^2+^ result in appearance of a new subpopulation of nucleosomes with a distribution centered at E_PR_=0.4 (Figure 1g). This subpopulation of N_P_ nucleosomes has significantly increased distance between DNA gyres near the location of fluorescent labels. The data show that Zn^2+^ ions specifically affect the interaction of PARP2 with nucleosomes.

### Interactions of PARP2 with nucleosomes in the presence of Zn^2+^ ions

According to spFRET data, similar effects of Zn^2+^ions on the structure of PARP2-nucleosome complexes are detected at Zn^2+^ concentrations ranging from 0.3 to 3 mM (Figure 2a). In the presence of Zn^2+^ ions, formation of complexes between PARP2 and nucleosomes begins at PARP2 concentration of *ca*. 12.5 nM, and 50% decrease in free nucleosomes occurs at 19±7 or 40±10 nM PARP2 according to spFRET microscopy or EMSA, respectively (Figure 2b, c). At the same time, a similar concentration dependence of PARP2 binding to nucleosomes was observed in Zn^2+^-free buffer A (Figure 1b). Thus, Zn^2+^ ions do not significantly affect the affinity of PARP2 for nucleosomes; however, Zn^2+^ ions induce a significant change in the structure of nucleosomes within the PARP2-nucleosome complexes.

**Figure 2.**
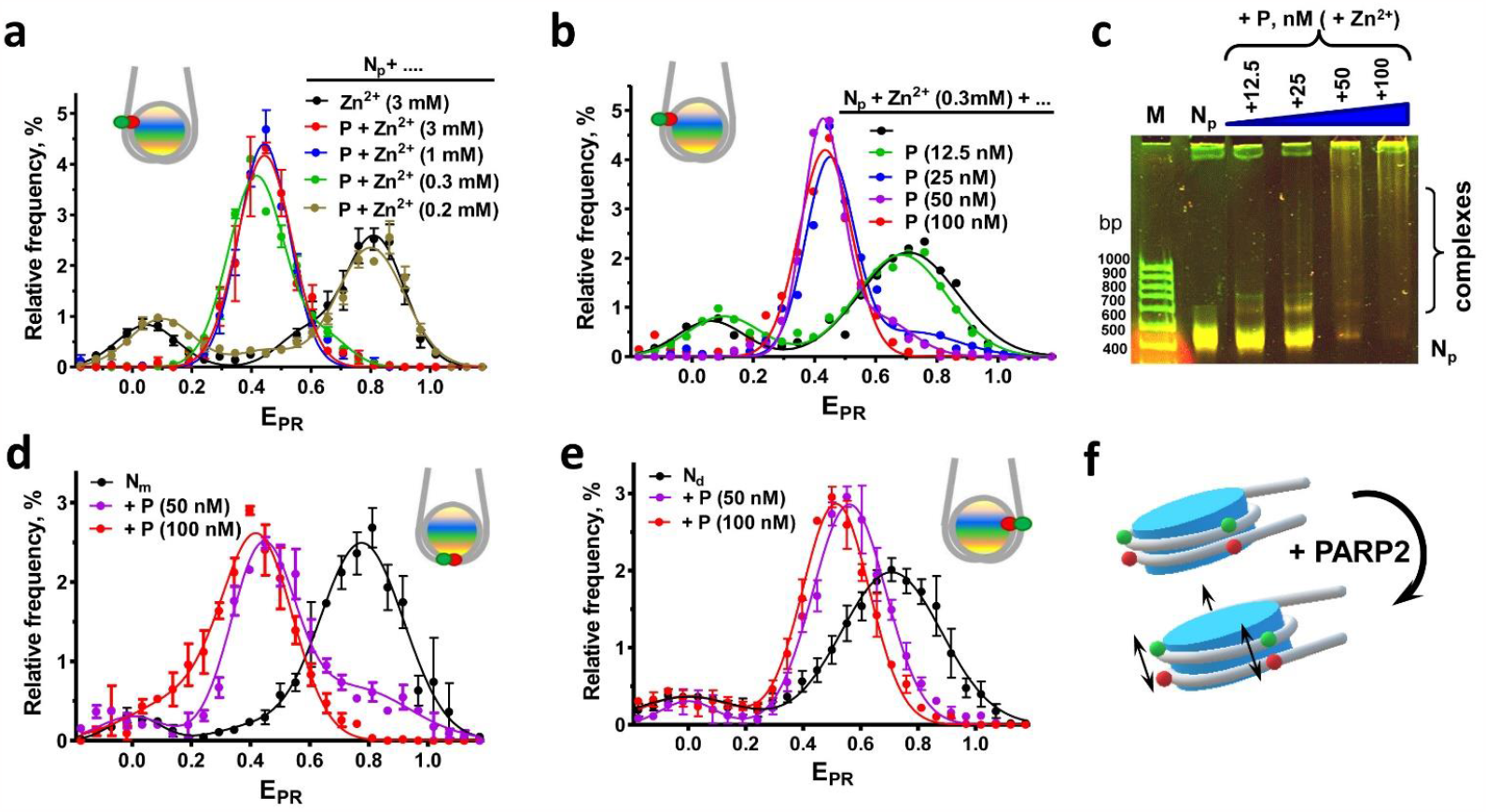
Zn^2+^-dependent nucleosome reorganization by PARP2. **a)** E_PR_ profiles of N_P_ nucleosomes and their complexes with PARP2 (P) in buffer A supplemented with different concentrations of Zn^2 +^ions. **b)** E_PR_ profiles of N_P_ nucleosomes mixed with different concentrations of PARP2 (P) in buffer A supplemented with 0.3 mM Zn^2+^ ions. Typical E_PR_ distributions of nucleosomes are shown (∼3000 particles per experiment). **c)** Analysis of N_P_ nucleosomes and their complexes with PARP2 (P, 12.5-100 nM) formed in buffer A supplemented with 0.3 mM Zn^2+^ ions by non-denaturing PAGE. M – DNA markers. **(d, e)** E_PR_ profiles of N_m_(**d**) and N_d_ (**e**) nucleosomes and their complexes with PARP2 (P) in buffer A supplemented with 0.3 mM Zn^2+^ ions. **f)** Proposed conformational changes in nucleosomal DNA induced by PARP2 binding. Green and red balls marks positions of the labels used to detect the conformational changes. Black arrows indicate an increase in the distance between DNA gyres detected using spFRET microscopy. **(a, d, e)** Statistics: mean±SEM; 3 independent experiments; ∼3000 particles per experiment.

Although two types of PARP2-nucleosome complexes are formed in the presence of Zn^2+^ ions according to EMSA (Figure S1a, b), the structures of these complexes are similar according to spFRET analysis (Figure 2b).

To characterize Zn^2+^-dependent structural changes induced by PARP2 within different regions of the nucleosomal DNA (the 35/112 and 57/135 bp regions), the N_m_and N_d_ nucleosomes (Figure 1a) were studied using spFRET microscopy (Figure 2d, e). As in the case of the N^P^ nucleosomes, spFRET microscopy in buffer A containing Zn^2+^ ions detect two subpopulations of particles for both N_m_ and N_d_ nucleosomes: a major subpopulation having higher E_PR_ values and a minor subpopulation having lower E_PR_ values (Figure 2d, e). Addition of Zn^2+^ ions and PARP2 to nucleosomes resulted in formation of a rather uniform structural state of nucleosomes having E_PR_ values centered at 0.4 and 0.5 for N_m_ and N_d_ nucleosomes, respectively (Figure 2d, e). These changes in the E_PR_ profiles of nucleosomes correspond to an increase in the distance between base pairs 35 and 112 as well as between base pairs 57 and 135 in the neighboring DNA gyres after formation of the complexes. Together with the data for N_P_ nucleosomes (Figure 2a, b), the obtained results show that the structural changes induced by PARP2 in the presence of Zn^2+^ ions affect the entire nucleosomal DNA (Figure 2f).

EDTA has a very high affinity for Zn^2+^ ions (*K*_*d*_∽10^−16^ M) (30, 31) and competitively removes Zn^2+^ from complexes with many proteins in a few minutes after their exposure to the chelator (31, 32). In contrast, addition of EDTA to the PARP2-nucleosome complexes preformed in the presence of Zn^2+^ does not induce changes in their structure even at the thirtyfold molar excess of EDTA over Zn^2+^ ions in solution (Figure 3a). These results allow us to propose that Zn^2+^ions involved in the formation of the PARP2-nucleosome complex are hidden inside this complex that hinders their extraction with EDTA.

**Figure 3.**
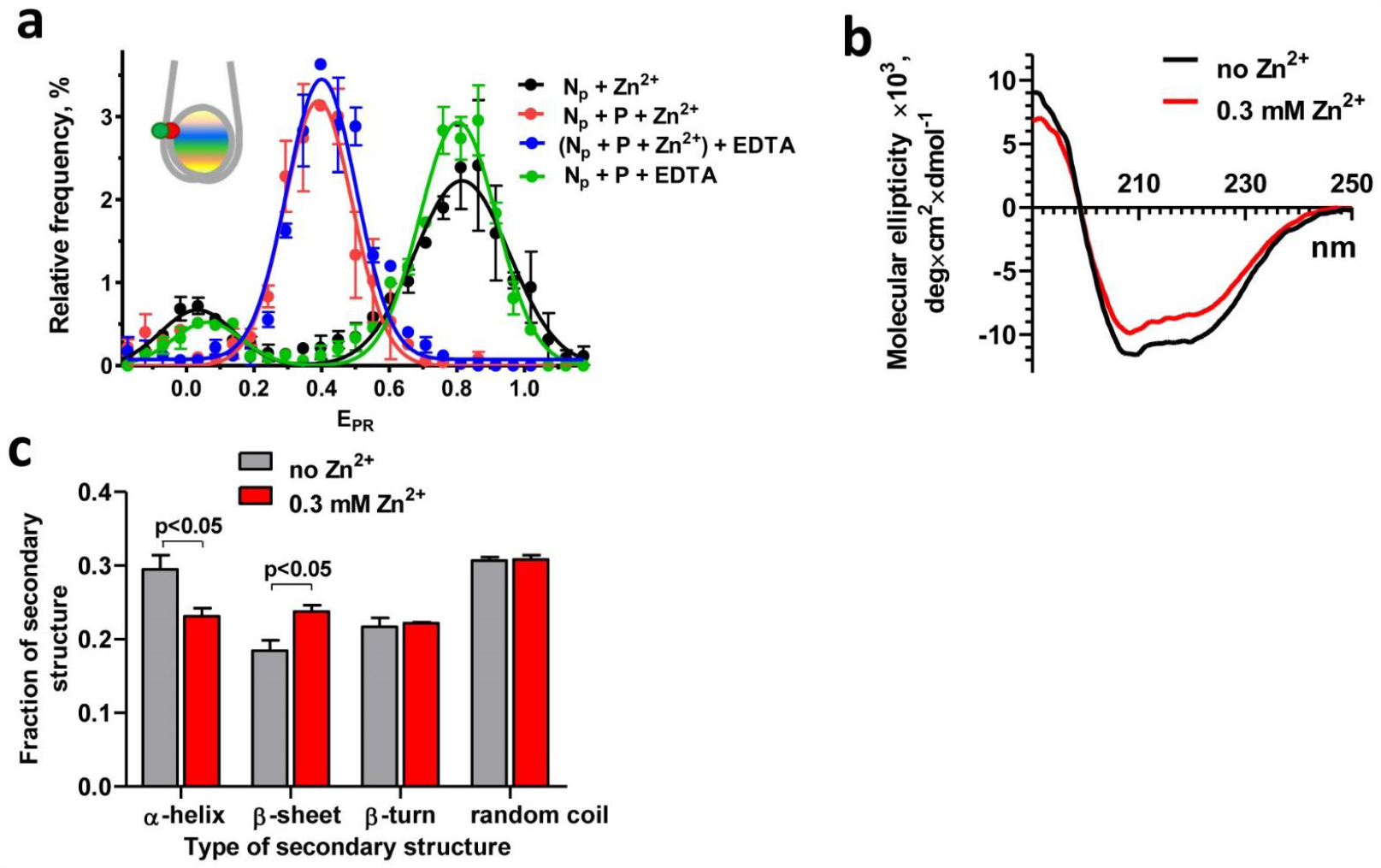
Analysis of Zn^2+^ interaction with PARP2 and PARP2-nucleosome complexes. **a**) Dependence of E_PR_ profiles of N_P_ nucleosomes and their complexes with PARP2 (P, 200 nM) on the presence of EDTA. Samples were measured in buffer A in the presence or absence of 0.3 mM Zn^2+^ ions. EDTA (10 mM) was added to the preformed PARP2 complexes with nucleosomes. Statistics: mean±SEM; 3 independent experiments; ∼3000 particles per experiment. **b)** CD spectra of PARP2 in the absence or presence of 0.3 mM Zn^2+^ ions. **c)** Content of canonical types of secondary structures in PARP2 according to the analysis of CD spectra of PARP2 measured in the absence or presence of 0.3 mM Zn^2 +^ions.

### Zn^2+^ binds to PARP2 and affects its secondary structure

Since no apparent Zn^2+^-binding domains were described for PARP2, Zn^2+^-dependent structural changes in a nucleosome induced by PARP2 binding (Figure 2) might indicate that Zn^2+^ ions specifically interact with PARP2, possibly altering conformation of the domain(s) of PARP2 that are responsible for the interaction with a nucleosome. To evaluate this possibility, circular dichroism (CD) spectra of PARP2 were measured before and after addition of 0.3 mM Zn^2 +^ions. The CD spectra differ noticeably (Figure 3b), detecting Zn^2+^-induced conformational changes in PARP2 that are related to a decrease in the α-helix content and a complementary increase in the content of β-sheet (Table S1, Figure 3c). Taking into account that the presence of Zn^2+^ does not affect the structure of intact nucleosomes (Figure 1e), the Zn^2+^-induced alterations in the conformation of PARP2 likely drive the changes in the mode of PARP2 interaction with a nucleosome and reorganization of the nucleosome structure in the complex.

### Effect of NAD+ on the structure of Zn^2+^-PARP2-nucleosome complexes

Addition of NAD^+^ substrate to the PARP2-nucleosome complexes activates a catalytic function of PARP2 and induces PARylation of neighboring proteins as well as autoPARylation of PARP2 (Figure 4a). Efficiency of autoPARylation of PARP2 depends on the concentration of added NAD+ (Figure 4a) and results in dissociation of PARP2 from the complexes with nucleosomes independently from the presence of Zn^2+^ ions (Figure 4b, c). The dissociation of the complexes likely occurs because of electrostatic repulsion between negatively charged PARylated PARP2 and nucleosomal DNA (1, 33, 34). According to the data of EMSA and spFRET microscopy, nucleosomes are restored their intact conformation after dissociation of PARP2 (Figure 4b, d). Thus, Zn^2+^-dependent structural changes induced in nucleosomes by PARP2 are reversible.

**Figure 4.**
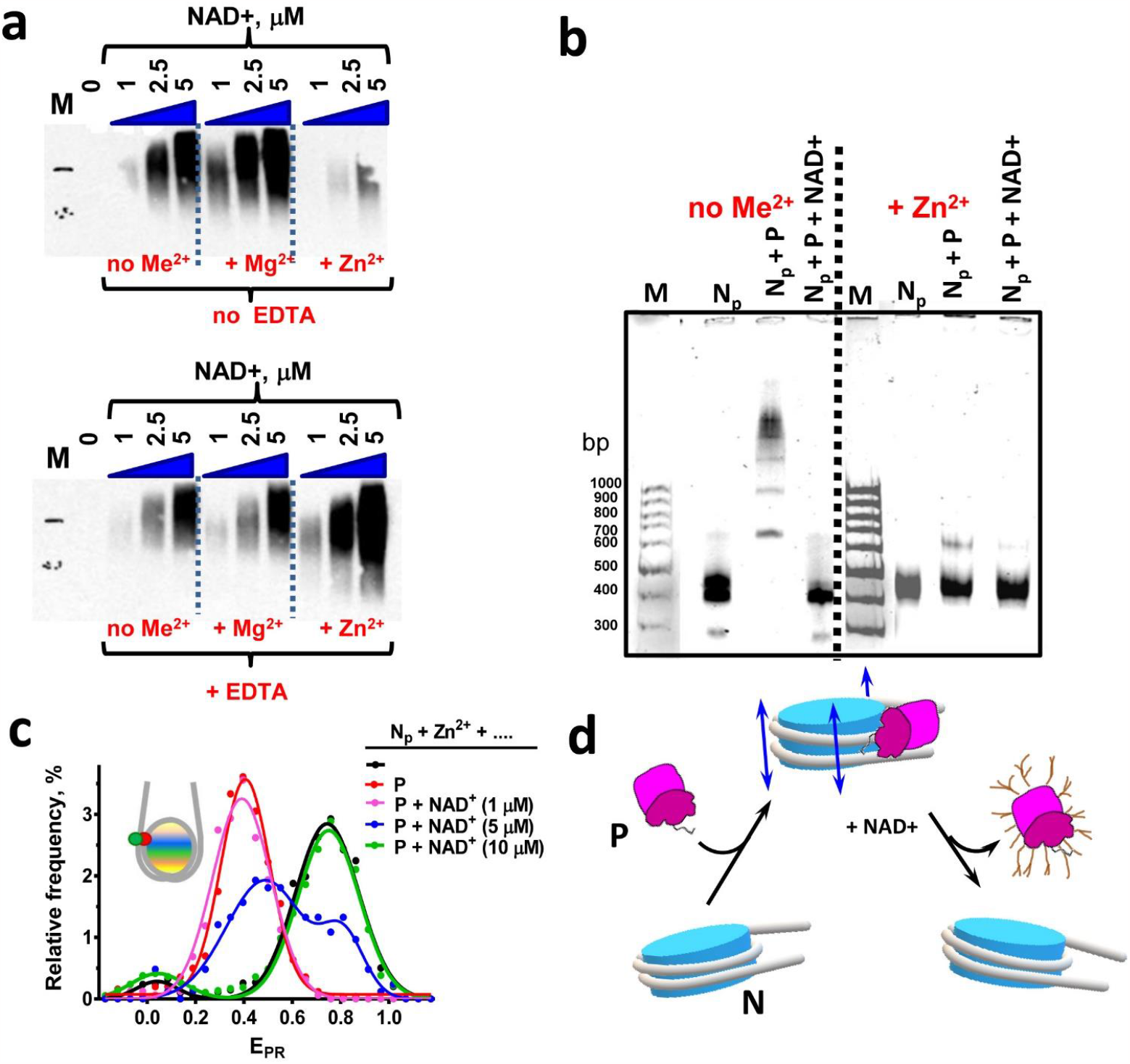
Enzymatic activity of PARP2 in the complexes with nucleosomes. **a)** WB analysis of NAD+-dependent PARylation by PARP2 in PARP2-nucleosome complexes in the presence and absence of EDTA (10 mM), Mg^2+^ (5 mM) and Zn^2+^ (0.3 mM). Me^2+^ - any divalent cations. b) Analysis of the complexes of N^p^ nucleosomes with PARP2 (100 nM) in the absence and in the presence of Zn^2+^ ions (0.3 mM) and NAD+ (10 µM). Me^2+^ - any divalent cations. **b)** Typical E_PR_ profiles of N_P_ nucleosomes and their complexes with PARP2 (P, 100 nM) after addition of different concentrations of NAD+. Samples were measured in buffer A supplemented with 0.3 mM Zn^2+^ ions. Statistics: ∼3000 particles per experiment. **c)** Changes in the structure of nucleosomes (N) in the presence of PARP2 (P) and NAD+. NAD+-meditated auto-PARylation of PARP2 results in dissociation of PARP2 from the nucleosomes, accompanied by recovery of the intact nucleosome structure.

The presence of Mg^2+^ or Zn^2+^ ions differentially affect the efficiency of PARP2-mediated PARylation: Mg^2+^ ions enhance, while Zn^2+^ ions decrease the efficiency of PARylation (Figure 4a). Addition of excess of EDTA, which chelates divalent ions, does not affect PARylation in the buffer without divalent ions and totally reverses the effect of Mg^2+^ ions (Figure 4a). Surprisingly, when Zn^2+^ ions are present in the buffer, addition of EDTA not only abolishes Zn^2+^-induced suppression of PARylation, but also strongly enhances the efficiency of PARylation (see Discussion).

## Discussion

PARP2 forms several complexes with a nucleosome, in which the conformation of nucleosomal DNA is not affected (Figure 1d, g). Formation of the complexes occurs both in the absence of divalent cations and in the presence of Mg^2+^ or Ca^2+^ions. A number of the complexes increases with the increase in concentration of PARP2: up to five complexes were detected by EMSA (Figure 1b, f). At least three of the complexes contain one nucleosome and likely contain different numbers of PARP2 molecules (Figure 1c). Binding of PARP2 to DNA occurs through the interactions of the N-terminal region and neighboring WGR domain of the enzyme (1, 15, 16). Taking into consideration the ability of PARP2 to recognize DNA double-strand breaks, two PARP2 molecules might be bound to the ends of nucleosomal DNA. At least one PARP2 molecule might be bound to the nucleosome core region, due to the higher affinity of PARP2 to nucleosomes as compared to DNA (13, 20). The presence of the additional complexes could be explained by the ability of PARP2 to form homodimers at the concentration higher than 50-80 nM with the apparent dissociation constants of 152 and 101 nM in the absence and presence of DNA, respectively (35). Assuming that this capability is preserved during the interaction with nucleosomes, formation of the additional PARP2-nucleosome complexes observed at 100-200 nM of PARP2 (Figure 1b) could be attributed to dimerization of PARP2 molecules bound to the nucleosome and/or to the ends of nucleosomal DNA.

Formation of PARP2-nucleosome complexes with different stoichiometry occurs nearly simultaneously (Figure 1b) indicating similar affinities of the components of these complexes to each other. Indeed, this complicates both detection of these complexes and measurements of their dissociation constants (*K*_*d*_) by many techniques. These complications most likely explain the failure to recognize formation of different PARP2-nucleosome complexes in earlier experiments (13, 20), and relatively high values of *K*_*d*_, which were reported previously (76 nM (20) and 150 nM (13)). According to our measurements, 50% of nucleosomes are involved in the formation of complexes with PARP2 at 40±10 nM PARP2 (Figure 1b). Considerably lower *K*_*d*_ (11 nM) was reported for the PARP2 complex with the 18 bp duplex oligonucleotides (20), while the *K*_*d*_ values for the complexes of PARP2 with the 28 bp duplex oligonucleotides depended on the type of the introduced DNA defect and varied from 37 to 178 nM (1, 15, 16). The *K*_*d*_ values of 11 and 37 nM probably describe the affinity of the 1:1 complexes, since direct binding of several PARP2 molecules to a short DNA fragment could be difficult due to possible steric interference between the bound PARP2 molecules, while dimerization of PARP2 occurs at much higher concentrations of PARP2 (35). Affinity of PARP2 for longer ds DNA (147 bp) was reported to be considerably lower, with *K*_*d*_ values variable in the range from 840 nM for intact DNA to 58 nM for DNA having a gap (13, 20).

The presence of submillimolar concentrations of Zn^2+^ ions in solution results in the PARP2-mediated structural reorganization of nucleosomal DNA (Figure 2) and affects autoPARylation activity of the enzyme (Figure 4a). These effects were most likely mediated by direct interaction of Zn^2+^ ions with PARP2, resulting in the local alteration of the secondary structure of the enzyme (Figure 3). The structural changes induced in nucleosomes by PARP2 in the presence of Zn^2+^ ions are almost completely reversible after dissociation of the autoPARylated enzyme.

Total concentration of zinc in cells is 0.2-0.3 mM (36), and 30-40% of this zinc is localized in the nucleus (37). Nuclear Zn^2+^ concentration increases during the S phase of a cell cycle compared to the G1 phase (38). Some processes (for example, nitrosative stress) induce considerable changes in Zn^2+^ concentration in the nucleus (39) or (more often) in cytoplasm (40, 41). In turn, changes in the concentration of cytoplasmic Zn^2+^ can induce quick response in the nuclear zinc concentration (42). Therefore, local transient changes in Zn^2+^ concentration within the nucleus could induce the change in the mode of interaction of PARP2 with nucleosomes and the autoPARylation activity of the enzyme.

After the Zn^2+^-mediated changes in the interaction mode of PARP2, any decrease in the Zn^2+^ concentration is not able to reverse these changes until dissociation of autoPARylated PARP2 occurs (Figure 3a), raising a question about the functional role of the Zn^2+^-mediated interactions of PARP2 with a nucleosome. The mechanism of DNA damage response (DDR) includes initial recognition of damaged sites by PARP1 and PARP2, activation of these enzymes in complexes with DNA, autoPARylation and PARylation of neighboring proteins including histones. Formation of PAR polymers attracts specialized DNA repair proteins to the damaged sites, where they substitute autoPARylated PARP1 and PARP2, which then dissociate from the complexes with DNA (43). It remains unclear why the damaged sites retain the increased affinity for DNA repair proteins after dissociation of PARP1 and PARP2, if before the binding of these enzymes, the sites were not recognized by the repair proteins. In the case of nucleosomes, one of the possible reasons is alteration of the structure of nucleosomal DNA induced by PARP1 (24) and by PARP2 in the presence of Zn^2+^ (Figure 2b). One possibility is that PARP-induced rearrangement of the nucleosome structure induces formation of the binding sites for some of the repair proteins. Beside DNA repair, PARP-induced reorganization of nucleosomes can have implications for such nuclear processes as transcription and replication, where the temporary partial reorganization of nucleosomes is required to facilitate propagation of transcription machine and replication fork, respectively.

According to our data, autoPARylation activity of PARP2 is modulated not only by Zn^2+^ ions, but also by Mg^2+^, and the ions can induce the opposite effects on PARP2 activity (Figure 4a). In accordance with the modern paradigm (1, 16, 43), autoPARylation activity determines the time of PARP2 association with a nucleosome (time of accumulation of a negative charge leading to PARP2 dissociation from a nucleosome), the rate of recruitment of the repair proteins and the dynamics of the early stages of DDR. Dependence of autoPARylation activity on the concentration of divalent cations was also reported for PARP1 (44). It seems that divalent cations are active players in DDR involving both PARP1 and PARP2, and transient changes in the concentration of the cations could modulate their contribution to the DDR dynamics. Futhermore, certain combinations of divalent cations were reported to synergistically potentiate autoPARylation activity of PARP1 (44).

To modulate the enzymatic activity of PARP2, the binding site for divalent cations should be localized at or very close to the catalytic domain of PARP2. Since EDTA was able to cancel the effect of Mg^2+^ on PARP2 activity in the complex with the nucleosome (Figure 4a), the binding site of Mg^2+^ is exposed to solution. EDTA was also able to change the effect of Zn^2+^ on the PARP2 activity from inhibiting to stimulating (Figure 4a), while it was not able to cancel the effect of Zn^2+^ on the structure of PARP2-nucleosome complex. Based on these data, we propose existence of two functionally different sites of Zn^2+^ binding in PARP2. One site is positioned at or close to the catalytic domain of PARP2 and is exposed to solution. This site is responsible for the negative regulation of PARP2 activity with Zn^2+^. The second site of Zn^2+^ binding is responsible for the altered mode of PARP2 binding to the nucleosome. It is most likely positioned at the interface of PARP2-nucleosome interaction and is hidden inside the complex and therefore inaccessible to EDTA. This binding site is also responsible for the enhancement of PARP2 activity, when Zn^2+^ is removed from the first binding site by EDTA (Figure 4a). Note that Zn^2+^ ions specifically affect the autoPARylation activity, but not the PARP2 affinity for the nucleosome.

In summary, autoPARylation activity of PARP2 is strongly modulated by divalent cations, and the effect of Zn^2+^ ions is more complex as compared to the effect of Mg^2+^ ions. Zn^2+^ ions are able to change the mode of PARP2 interaction with the nucleosome that in turn could modulate the chromatin-binding and DNA damage recognition functions of PARP2. Our data show that even in the absence of specialized Zn^2+^-binding domains like zinc fingers, PARP2 is Zn^2+^-dependent protein, and the presence of Zn^2+^ modulates PARP2 activity and its interaction with nucleosomes.

## Acknowledgments

We thank M. Kutuzov and O. Lavrik (Institute of Chemical Biology and Fundamental Medicine, Novosibirsk, Russia) for a gift of PARP2. We thank N. Gerasimova, G Armeev, D. Koshkina for discussion.

