## Supplemental Data for "Zinc-dependent Nucleosome Reorganization by PARP2"

**Table S1. Content of canonical types of secondary structures in PARP2 according to the analysis of CD spectra of PARP2 measured in the presence or absence of $Zn^{2+}$.**

| Structure | no $Zn^{2+}$ | | 0.3 mM $Zn^{2+}$ | | P-value |
|---|---|---|---|---|---|
| | Mean | SEM | Mean | SEM | |
| α - helix | 0.295 | 0.011 | 0.231 | 0.007 | 0.024 |
| β-sheet | 0.184 | 0.008 | 0.237 | 0.005 | 0.022 |
| β-turn | 0.217 | 0.007 | 0.222 | 0.001 | 0.765 |
| random coil | 0.307 | 0.003 | 0.308 | 0.004 | 0.828 |

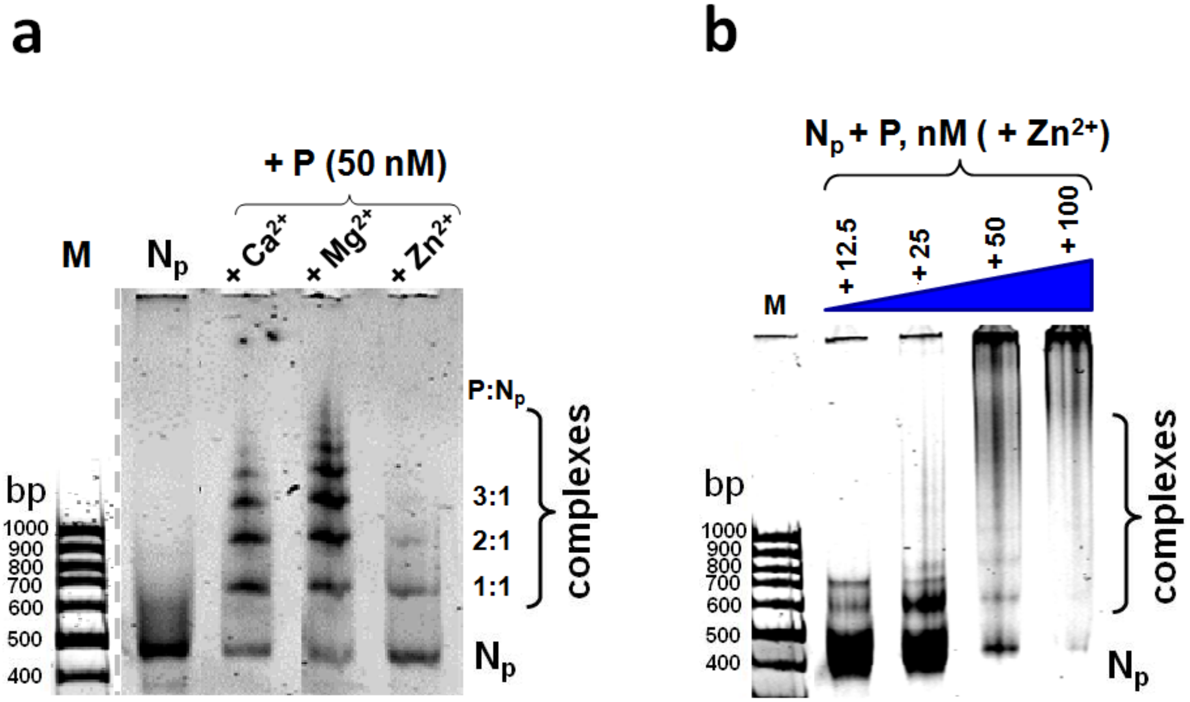


**Figure S1. PARP2 forms complexes with nucleosomes of different stoichiometry.**

**a)** Analysis of $N_P$ nucleosomes and their complexes with PARP2 (P, 50 nM) formed in buffer A in the absence of divalent cations ($N_p$) or in the presence of $Zn^{2+}$ (0.3 mM), $Ca^{2+}$ (5 mM) or $Mg^{2+}$ (5 mM) ions by non-denaturing PAGE. P: $N_P$ is a proposed stoichiometry of the PARP2 complexes with nucleosomes observed in the gel. M – DNA markers.

**b)** Analysis of $N_P$ nucleosomes and their complexes with PARP2 (P, 12.5-100 nM) formed in buffer A supplemented with 0.3 mM $Zn^{2+}$ ions by non-denaturing PAGE. M – markers.
